# TxCyto: A machine learning framework for estimating cytokine activity from whole transcriptome

**DOI:** 10.64898/2026.09.28.754405

**Authors:** Rahul Kumar, Shan Li, Peng Jiang, Sridhar Hannenhalli

## Abstract

Cytokines are critical mediators of intercellular communication, and a comprehensive characterization of their activity is essential for understanding health and disease. Existing tools to infer cytokine activity rely on experimental measurements. However, such measurements are available only for a small minority (43) of cytokines, and moreover, cytokine activity and response are highly context-specific, making a comprehensive experimental profiling across tissues, disease states, and biological contexts impractical. To address this gap, we developed TxCyto - a deep learning-based framework that infers the activity of cytokines, and more broadly of the tumor secretome, directly from the whole transcriptome profile of a sample. Trained on pan-cancer TCGA tumor transcriptomes, TxCyto was extensively validated in multiple independent datasets, including cytokine perturbation experiments. Across multiple cancer immunotherapy cohorts, TxCyto identified cytokines whose predicted activity was associated with therapeutic response. Furthermore, in spatial transcriptomic data for Liver cancer, TxCyto discovered spatial niches associated with response to immunotherapy. Overall, we develop a machine learning tool-TxCyto, for predicting the activity of 645 cytokines and tumor secretome from readily available whole transcriptomes. The TxCyto framework is generally applicable to other classes of regulatory molecules and TxCyto code base, and the tools are provided at https://github.com/Rahulncbs/TxCyto.

## Introduction

Cytokines are a diverse family of secreted signaling proteins that are primarily involved in immune activity, but more broadly, they regulate cellular behavior in multicellular organisms [1,2]. Cytokines are key biomarkers in a variety of disease conditions such as infection, inflammation, autoimmunity, and cancer [3,4], as well as therapeutic targets for various indications such as rheumatoid arthritis (TNF/IL-6) [5], psoriasis (IL-17/IL-23) [6,7], inflammatory bowel disease (IL-23/TNF) [8], and cancer immunotherapy (IFN-γ) [9,10]. Cytokines are typically transiently produced at low level [11]. Furthermore, they exhibit pleiotropy, redundancy, interactions, and multiple modes of action (autocrine, paracrine, endocrine) [12]. These properties of cytokines make them highly challenging to profile and functionally characterize. Given their profound clinical importance, knowing the downstream transcriptomic mediators of cytokines, as well as ability to estimate cytokine activity from transcriptomic data is highly important.

CytoSig has comprehensively compiled from the literature 20,591 nonredundant individual transcriptomics profiles for cytokine, chemokine and growth hormone treatment experiments in 962 diverse human cellular contexts covering 43 cytokines [13]. Learning from differential gene expression upon cytokine-specific perturbations, independent of cell type context, CytoSig builds a machine learning tool to predict cytokine activities from a given transcriptomic profile. A major limitation, however, is the relatively small number of cytokines (43 out of hundreds of known human cytokines [14]) subjected to controlled perturbation experiments. Equally importantly, cytokines are perturbed only in a few immune cell types under specific conditions which may not sufficiently represent physiological diversity in which they function in vivo [15]. It is challenging to experimentally profile each cytokine comprehensively in various physiologically relevant contexts, and therefore, a computational model which can learn from existing data and reliably predict the cytokine activity in multiple contexts, across a much larger number of cytokines, is warranted.

Tumor microenvironment (TME) is composed of different cell types, most of which contribute to the global cytokine pool in the TME, and most cell types respond to the available cytokines in the environment, regardless of the producing cell, and respond to it in a manner that is reflected in their transcriptome [16,17]. The key premise then is that the global transcriptome across all cell types may reflect the overall cytokine in the TME and thus can be used to estimate cytokine activity. This premise applies not only to cytokines but extends to all secreted molecules sensed by cells [18]. While we use the term ‘cytokine’ for simplicity, we have included a much broader collection of 645 proteins secreted in the TME compiled from the reference database [19]. While our goal is to estimate cytokine activity and protein level of cytokine is a better proxy for its activity, due to limited availability of cytokine protein quantification, we use the global RNA level of cytokine as a proxy for its activity. Based on the above premise, here we developed a deep learning tool - TxCyto, trained on bulk TCGA data, to predict cytokine expression from the transcriptome of a sample.

We validated TxCyto in several independent data sets, including cytokine blocking and cytokine induction experiments. We show that TxCyto prediction accuracy is maximal when trained on bulk TCGA data, compared to both the model trained on deconvolved cell type-specific transcriptome from bulk, as well as the model trained on a large cohort of scRNA-seq data. We applied TxCyto to pre-treatment transcriptomic data of immunotherapy cohorts spanning five tumor types -- Breast cancer, Melanoma, Head and neck cancer, Non-small cell lung cancer, and clear cell renal cell carcinoma. The model consistently reveals some of the known cytokine-axis (IFNγ and T-cells recruiting chemokines such as CXCL9, CCL10) of favorable immunotherapy responses across multiple cancer types. Furthermore, it also reveals the tissue-specific nature of immunotherapy resistance and the associated cytokines. We have provided tissue-specific cytokines that could be tested for their potential role as a biomarker in stratifying patients by potential immunotherapy response. Next, we applied TxCyto on spatial transcriptomics data related to ovarian cancer and hepatocellular carcinoma (HCC) treated with neoadjuvant chemotherapy and novel immunotherapy combination therapies respectively. Using TxCyto predicted cytokine activity level, we were able to classify the spatial niches and donors into responder and non-responder achieving AUROC values of 0.77 and 0.75 respectively. Overall, we provide a general cytokine activity prediction tool based on transcriptomic data, which is broadly applicable, and substantially expanding the scope of current resources for cytokine activity prediction. We provide the pre-trained TxCyto tool as an easy-to-use open-source tool at https://github.com/Rahulncbs/TxCyto.

## Results

### TxCyto accurately estimates cytokine activity across cytokines and tissue contexts

Figure 1 illustrates the TxCyto pipeline. TxCyto uses a fully connected neural network (FCNN)-based deep learning model that learns the relationship between the global transcriptomic state and cytokine activity in a sample. TxCyto is based on the premise that (i) the cytokine activity in a tissue sample is reflected in the transcriptomic states of the constituent cells in the tissue, (ii) cytokine activity estimate based on the global transcriptome is more robust to technical and biological noise associated with direct measurement of cytokine, and is more sensitive to the biological context. TxCyto was initially trained and tested via 5-fold cross-validations on the pancancer TCGA bulk transcriptomic data spanning 33 cancer types and 9491 samples and validated in multiple independent cohorts (See Material and Methods).

**Figure1:**
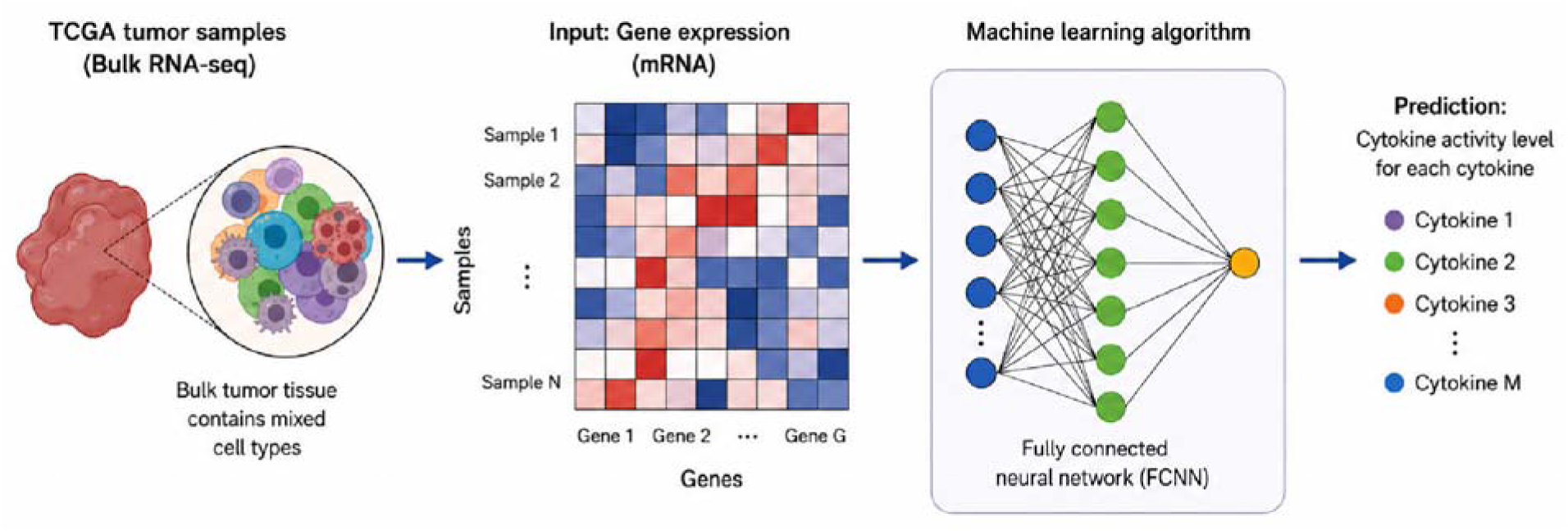
Schematic of the TxCyto pipeline to predict cytokine activity: The model is based on a fully connected neural network (FCNN). It takes TCGA pan cancer bulk transcriptomic data as input to train and predicts cytokine activity in a tissue sample.

We used 6075 common genes as the input features to the model to independently predict the activity of each of the 645 different cytokines (Supplementary Table 1). The cross-validation (CV) accuracy is computed using Pearson’s correlation between predicted cytokine activity level with actual mRNA expression level across the test samples. Besides the model based on bulk mRNA, we performed sample-specific deconvolution using CODEFACS [20] (Methods) to infer sample-specific expression of 11 cell types and additionally evaluated 11 cell type-specific models (Methods), i.e., given the cell type-specific expression in a tissue how well the model could predict the global cytokine level in the TME. The average CV accuracies of all 645 cytokines across 12 models (one bulk and 11 cell type-specific models) are shown in (Figure 2A). We found that the median CV accuracy across cytokine from the bulk transcriptome-based model is 0.78, higher than all other models; malignant cells-based model accuracy was close second. The bulk transcriptome and the malignant cells-based models have the most number of cytokines (482 and 442 respectively out of total 645 cytokines) with CV accuracy > 0.6 (Figure 2B). These results are consistent with the fact that the malignant cells dominate tumor biopsies (median purity score is ∼0.64 TCGA, Supplementary Figure 1). The results also suggest that the transcriptomes of other microenvironmental cell types do not capture the cytokine activity in the TME as good as the model based on malignant cells or overall bulk-transcriptome.

**Figure 2.**
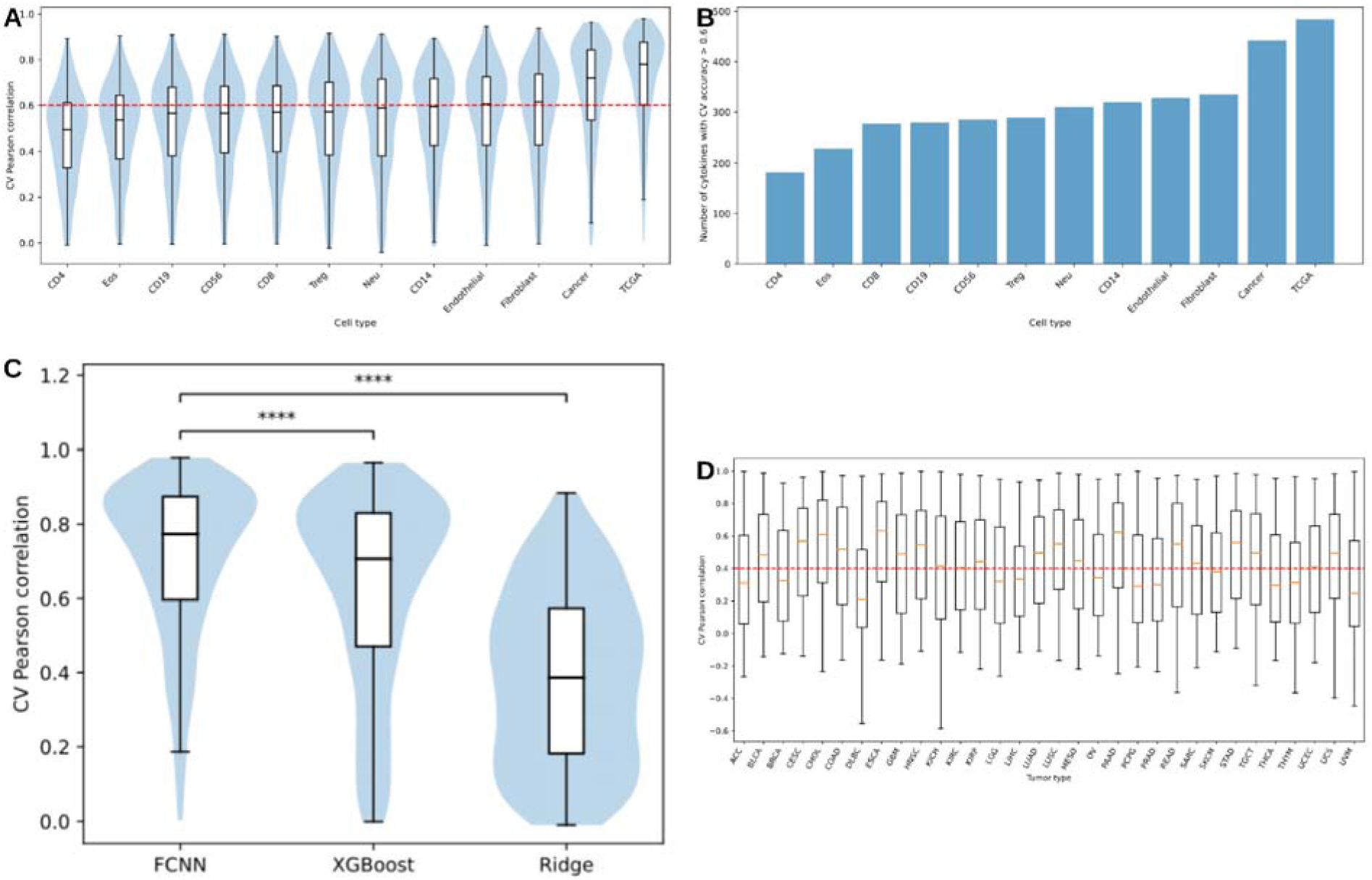
A) The distribution of the cross-validation (CV) accuracies of cytokines across 12 models based on deconvoluted cell type-specific transcriptome and TCGA bulk transcriptome. B) Plot showing the number of cytokines with CV accuracy>0.6 from each of the 12 models. C) Comparing the distribution of CV accuracy of the cytokine prediction from models based on FCNN, XGBoost and linear ridge regression. D) Evaluating the generalizability of the model based on TCGA bulk transcriptome using leave-one-tissue-out strategy, the x-axis represents the tumor type held out from training and used for evaluation. Y-axis is accuracy (Pearson correlation) between predicted and actual cytokine gene expression in the held out tissue.

Further, to assess relative advantage of the FCNN based non-linear architecture for this task, we compared its performance with those for a linear ridge regression-based model and XGBoost -- a nonlinear tree-based machine learning approach. We trained and validated all three model architectures on identical features and sample cohorts (see Methods). The FCNN model achieved substantially higher CV accuracy (median CV =0.78) across cytokines compared to ridge regression-based model (median CV =0.38), supporting the need for a non-linear model (Figure 2C). Even compared to a non-linear XGBoost model (median CV ∼0.70), FCNN showed small but significant improvement, highlighting the advantage of the FCNN architecture for the task (Figure 2C).

Next, we evaluated performance across all 33 TCGA tumor tissues using a leave-one-tissue-out strategy, in which each tumor type was held out from model training and used only for evaluation. Although predictive performance varies across tumor types, 19 out of 33 tissue types achieved median CV accuracy >0.4, supporting the generalizability of the model across diverse tissue contexts (Figure 2D). Based on the various comparisons above, for all downstream validations and applications, we use the FCNN model based on the TCGA pan-cancer bulk transcriptome.

Given that there is no single assay which can directly and comprehensively measure the cytokine activity at scale, in the following sections we sought to establish the biological validity of TxCyto predicted cytokine activity with several independent and complementary lines of evidence. These include direct protein quantification, causal perturbation experiments and orthogonal transcriptomic signature which includes downstream pathways and receptor expression concordance with the predicted cytokine activity.

### Tissue-level bulk transcriptomes capture coordinated multicellular cytokine signaling more robustly than pseudobulked single-cell profiles

The TME can be characterized using either bulk RNA sequencing or single-cell transcriptomic profiling. While scRNA-seq provides cellular compartment-specific transcriptome, it suffers from relatively smaller cohort sizes as well as variable dissociation efficiency across various cell types distorting their representations [21]. We nevertheless assessed the relative advantage of training a cytokine model based on scRNA-seq data, on a cohort of 1062 scRNA-seq samples, comprising all major cell types [22] (see Methods). As in the bulk data, epithelial/malignant cells were the most abundant cell type in the scRNA-seq data (Supplementary Figure 2). We generated pseudobulk transcriptomic profiles to approximate the bulk transcriptome of the tissue sample. We first assessed the relative advantage of using all cell types versus only the epithelial/malignant cell compartment to infer cytokine level in the sample. As shown in Figure 3A, combining all cell types provides a more accurate model than using only the epithelial/malignant compartment, in both the TCGA and scRNA-seq cohort. Furthermore, the TCGA-based model achieves higher cross-validation accuracy than the scRNA-seq-based model. Note that the above comparison addresses which transcriptomic representation best predicts tissue-level cytokine activity, not whether individual cell types respond differently to cytokine exposure — the latter is well established and does not conflict with our finding that coordinated, multicellular signal is most informative for cytokine activity inference.

**Figure 3.**
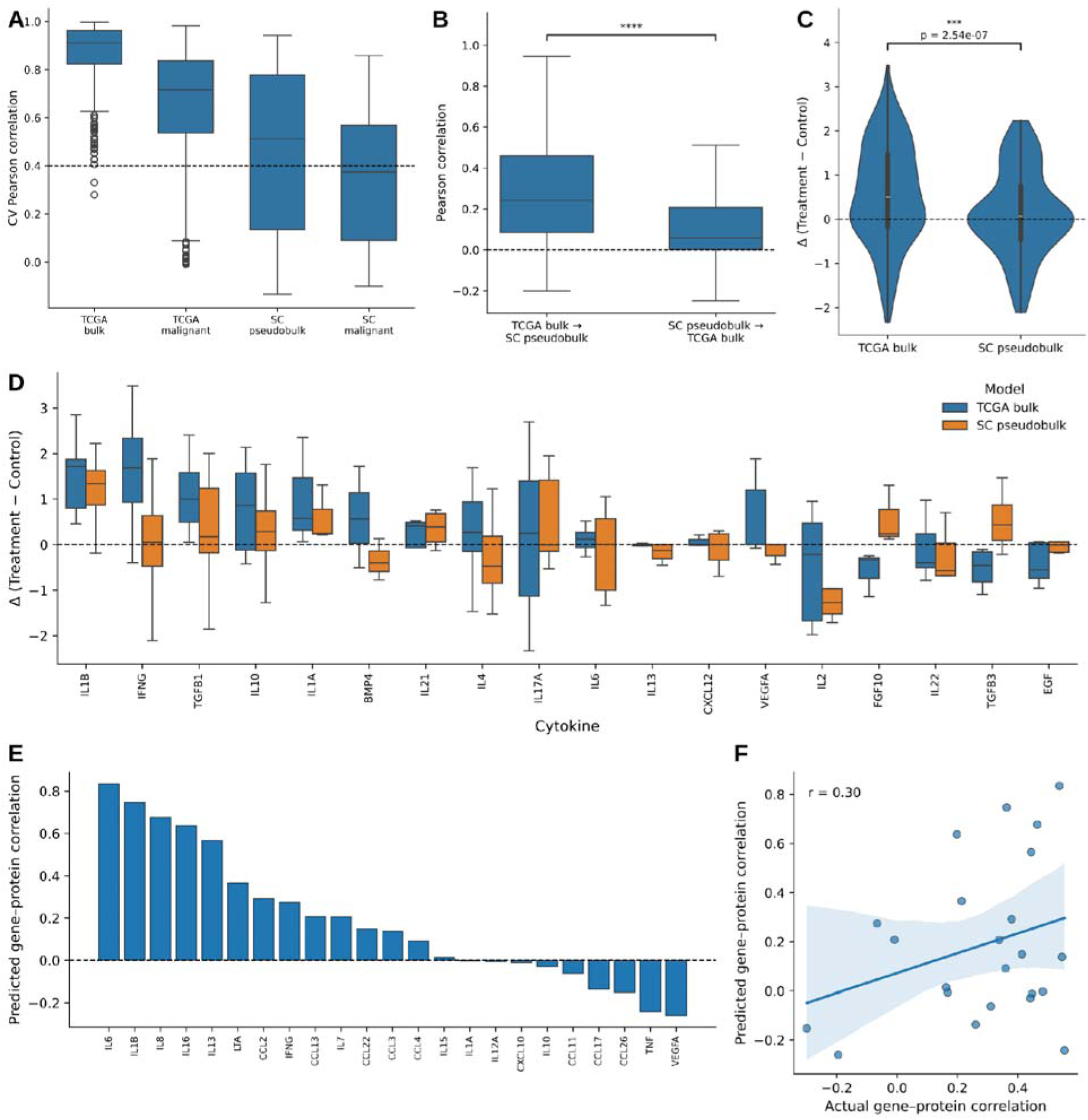
A) Comparing the TCGA bulk transcriptome-based model CV accuracy with single cell pseudo-bulk transcriptome-based model CV accuracy. The X-axis shows four different transcriptome-based models: TCGA bulk transcriptome, TCGA malignant cell-based transcriptome, Single-cell pseudobulk transcriptome, Single-cell (malignant only) pseudobulk transcriptome. Y-axis is CV accuracy. B) Comparing the Delta (treatment-control) across cytokine and cell types for the two models: (i) TCGA bulk transcriptome and (ii) single cell pseudobulk transcriptome. Here the cytokine is induced in the treatment condition, and the activity is compared with the control untreated sample. C) Cross-comparing the prediction accuracy of the model trained on TCGA bulk and tested on single cell pseudobulk and vice versa. D) Same as (C) but split by individual cytokines separately. E) Plot showing the Pearson correlation between predicted cytokine level with actual protein level (Y axis) in an Ovarian cancer cohort for individual cytokines (X axis). F) Scatter plot comparing the Pearson correlation between actual cytokine mRNA and the protein level (X axis) the Pearson correlation between predicted cytokine mRNA and the protein level (Y axis) in ovarian cancer.

Next, to more directly assess the relative advantage of the two approaches, we compared the accuracy of (1) the model trained on TCGA and tested on scRNA-seq pseudobulked cohort, with (ii) the model trained on scRNA-seq pseudobulked cohort and tested on TCGA. An identical set of input features were used in both analyses (see Methods). As shown in Figure 3B, the model trained on TCGA and tested on scRNA-seq pseudobulk is far more accurate than the other way round, suggesting that the TCGA bulk transcriptome better reflects the cytokine activity in the sample. Finally, we evaluated both the TCGA-trained and the scRNA-seq pseudobulk-trained models in an independent experimental dataset (see Methods, cytokine and its GEO accession ID is provided in Supplementary Table 2). A cytokine model is expected to infer a higher level of cytokine post-induction. We therefore evaluated the pre-induction to post-induction increase (cytokine response delta) in the inferred cytokine activity as per the model (trained on TCGA or scRNA-seq pseudobulk cohort). Figure 3C shows the distribution of cytokine response deltas inferred by the two models. Of the total 202 cytokine-experiment conditions compared, in 134 cases (∼66%) the TCGA-trained model produces delta (treatment-control) positive compared to 106 cases (∼52%), where scRNA-seq pseudobulk trained model produced positive delta. Figure 3D further breaks down the delta distributions in a cytokine-specific manner. A higher delta value indicates a greater ability of the model to distinguish cytokine-treated samples from control samples. Of the 18 cytokines examined, the median delta was positive for 10 cytokines when TCGA-trained model was used and for 9 cytokines when scRNA-seq pseudobulk-trained model was used. Collectively, these findings indicate that tissue-level bulk transcriptomes may preserve the coordinated multicellular cytokine signaling programs more robustly than dissociated single-cell transcriptomic representations.

Next, we evaluated the TxCyto predicted cytokine activity with respect to measured cytokine protein levels. Cytokine protein level is a better proxy of its activity than the RNA level, however, cytokines normally occur at low levels with short half-life [23], and there is a paucity of matched mRNA-protein level profiling making it difficult to directly model protein levels of cytokines from the sample transcriptome. Nevertheless, we evaluated our RNA-based TxCyto model on an independent cohort of 32 transcriptomic samples with matched quantification of 23 cytokines in ovarian cancer (Figure 3E). As expected, the model accuracy is higher for the cytokines with higher mRNA-protein correlations (Figure 3F), but interestingly, for 9 cytokines, the predicted cytokine expression is better correlated with the protein level than the observed expression is, suggesting that the model may better capture cytokine activity in some cases; we further assess this possibility below. Together, these results suggest that exploiting the bulk transcriptome, TxCyto effectively captures the activity of a large fraction of cytokines. Importantly, previous standard to estimate cytokine activity relies on perturbation data, which are very limited both in terms of number of cytokines profiled as well as in the tissue contexts; for instance, CytoSig [13] offers predictive models of 43 cytokines. In contrast, because TxCyto does not rely on cytokine perturbation experiments, it substantially expands the repertoire of cytokines whose activity can be estimated in a transcriptomic sample.

### TxCyto correctly predicts lower cytokine activity upon treatment with cytokine inhibitors

Above we validated TxCyto on cytokine induction experiments. Here, we assessed the extent to which TxCyto predicts a decrease in cytokine activity in samples treated with a cytokine inhibitor. We compiled transcriptomic datasets from multiple cytokine-blocking studies from the GEO database spanning diverse human physiological conditions such as lupus, psoriasis, atopic dermatitis, and rheumatoid arthritis synovium, in which patients were treated with cytokine inhibitors and RNA-seq profiles were generated from the lesional and non-lesional skin biopsy, whole blood samples pre and post treatment or at different time points and dosage. (details of the studies are provided in the Supplementary Table 3).

Cytokine activity inferred by TxCyto in the pre-treatment and the post-treatment transcriptomic samples were compared across these cohorts (see Methods for grouping the samples in different experimental contexts). As shown in Figure 4A, the TxCyto inferred delta is negative for the majority of cases (16 out of 20 (80% cases)) suggesting the cytokine activity decreases post treatment with cytokine inhibitors. The concordance between TxCyto and CytoSig was evaluated based on the consistency in the direction of delta. As shown in Figure 4B, both the methods inferred a negative delta in 11 out of 20 cases, whereas they disagreed on the direction of delta in 9 cases. Across the independent validation cohort treated with cytokine inhibitors, TxCyto predicts a negative treatment induced change in 16 of 20 cases compared to 15 of 20 cases from CytoSig, suggesting TxCyto provides a modest improvement in estimating cytokine activity across cohorts Figure 4C. Overall, despite the fact that TxCyto is trained on cytokine RNA expression, TxCyto can infer the expected decrease in cytokine activity upon treatment with cytokine inhibitors.

**Figure 4.**
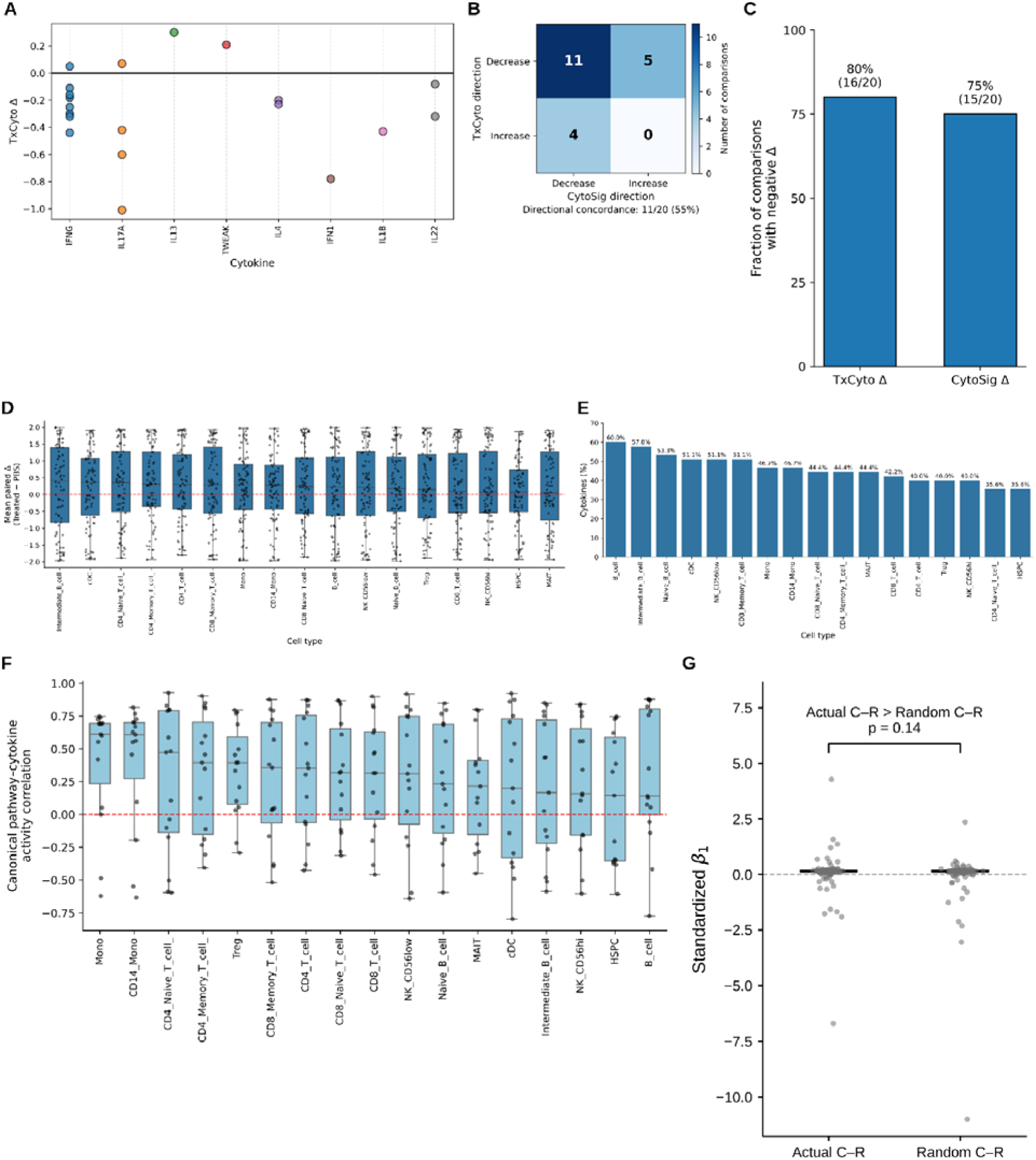
A) Plot showing the delta (treatment-control) inferred from TxCyto for different cytokines in sample cohorts treated with cytokine inhibitors and its control set. B) The heat plot showing the concordance between the sign of delta inferred from TxCyto and CytoSig. C) Comparing the fraction of cytokines with negative delta values inferred by TxCyto with that for CytoSig in sample cohorts treated with cytokine inhibitors. D) The plot showing the distribution of delta in different immune cell populations induced with cytokine and control set. Each cell type was pseudo bulked prior to application of TxCyto. E) The bar plot shows the fraction of cytokine significantly elevated in the treated sample compared to the control sample across 17 immune cell populations of PBMCs. F) The plot showing the distribution of the Pearson correlation values for 15 cytokine activity inferred from TxCyto and its downstream canonical pathway pairs across different immune cell populations. G) The plot showing the distribution β1 value for the matched cytokine– receptor pairs with that of randomized cytokine-receptor pairs. The β1 is the coefficient of receptor expression level.

### TxCyto predicts cytokine-stimulation across immune cell populations

Following up on TxCyto validation in cytokine-blocking experiments, here we validated TxCyto on one of the largest cohorts of cytokine-stimulated single-cell transcriptomic data, consisting of cell type–resolved scRNA-seq generated from human peripheral blood mononuclear cells (PBMCs) obtained from 12 donors and stimulated in vitro with 90 different cytokines (see Methods)[15]. We focussed on 45 cytokines with CV accuracy > 0.6 (see Methods). A total of 24 distinct immune cell populations were captured; 7 cell types were excluded which don’t have all 12 donors scRNAseq data present. After preprocessing, filtering for the number of samples and cytokines (see Methods), 45 cytokines across 17 immune cell types in treatment and control conditions were analyzed. For each type, single-cell RNA-seq profiles were pseudobulked, followed by cytokine activity prediction using the TxCyto before and after treatment. We first assessed the extent to which TxCyto captures the expected increase in activity of these 45 cytokines in each cell type upon treatment. As shown in Figure 4D, the median delta value (treatment minus control) was positive in the majority of cell types, suggesting that TxCyto captures the expected increase in cytokine-associated activity following cytokine treatment, again, despite being trained on cytokine RNA level in bulk data. Figure 4E shows the fraction of cytokines that were predicted to be significantly elevated (Wilcoxon test p-value <= 0.05) in treated samples relative to control samples in a cell-type specific manner. In several immune cell compartments such as B cells, CD8 memory T cells, dendritic cells, more than 50% of the cytokine exhibit significant increase in activity post-treatment, suggesting stronger treatment-associated cytokine activation programs in these cell types. Of the total 765 tests (45 cytokines x 17 cell types), 46% were significant at p value <= 0.05 (i.e., > 9-fold enrichment). The heatmap shown in the Supplementary Figure 3 shows the cytokines significantly elevated post treatment across different immune cell compartments, highlighting the variable responses of cytokine treatment across different immune cell populations. Given that TxCyto is trained on bulk transcriptome, and thus inherently relies on combined responses in multiple cell compartments, and therefore the transcriptomic changes in a specific cellular compartment may not sufficiently reflect the global cytokine shift, these results are encouraging. Together, these observations support TxCyto’s ability to capture cytokine induction in individual cell types, despite being trained on bulk transcriptome data, while revealing heterogeneous responses across immune cell compartments.

### Cytokine-specific downstream pathways positively correlate with predicted cytokine activity

Next, in the same cytokine-stimulated data-cohort used above, we further assessed whether the inferred activity of a cytokine correlated with its canonical downstream pathway score, computed independently using GSEA pathway enrichment method [24]. We literature-curated 15 cytokines along with their canonical downstream pathways provided in the Supplementary Table 4 [25,26]. For each cell type, we calculated the correlation between the TxCyto-inferred cytokine activity and their corresponding downstream pathway activity scores from GSEA in the treated samples. Figure 4F shows the distribution of the correlation values for the 15 cytokine-pathway pairs across different cell types. We specifically examined some of the key cytokines, including IFN-gamma, IFN-lambda1, IL-2, and IL-6 together with their established downstream pathways to assess the consistency of these associations across immune cell populations. Across the majority of immune cell types, the median correlation score remained consistently positive, suggesting that the cytokine activities inferred by TxCyto are concordant with the expected activation of their expected downstream signaling programs and likely reflect the true cytokine activity state of the cells.

### Predicted cytokine activity captures biologically relevant cytokine-receptor signaling relationship

Presence of cytokine receptor is required for a cytokine’s downstream effect on the transcriptome, which is used by TxCyto to estimate cytokine activity. We therefore tested the extent to which this expectation holds true based on 88 curated ligand–receptor pairs (Supplementary table 5). We used a human lung tumor bulk transcriptomic data cohort comprising 29 samples to examine the association between predicted cytokine activity and the corresponding receptor expression levels [27]. We hypothesized that known cytokine–receptor pairs would show stronger associations than randomized cytokine-receptor pairs. We fitted a regression model to estimate predicted cytokine activity in terms of (i) the observed cytokine expression level, and (ii) the observed receptor expression level. We then compared the receptor-associated regression coefficient β1 between real and random pairs (see Methods). As shown in Figure 4G, the median β1 value was modestly higher for the matched cytokine– receptor pairs than for the randomized pairs, suggesting that the predicted cytokine activity captures biologically meaningful receptor-mediated downstream signaling relationships.

### Identification of cytokines associated with immunotherapy response in multiple cancer types

Having validated the TxCyto in independent datasets, we applied it to multiple clinical immunotherapy cohorts to systematically evaluate the relationship between pre-treatment cytokine activity and immunotherapy response. The data included 19 studies, across 5 cancer types (Supplementary table 6). The detailed number of samples for each cancer type along with responder and non-responder labels is provided in the Supplementary table 7. Applying pre-trained TxCyto to these independent immunotherapy cohorts, we identified cytokines significantly associated with either favorable or unfavorable immunotherapy responses.

Cytokines were prioritized based on i) Effect size, defined as the difference in the mean predicted cytokine activity between responders and non-responders. and ii) Statistical significance, assessed using one-sided Mann-Whitney U test. Separate tests were performed to identify cytokines with significantly higher activity in responder (effect size>0) and significantly higher activity in non-responders (effect size<0). Further, P values were adjusted for multiple testing using the Benjamini–Hochberg procedure, and cytokines with FDR<0.25 were considered significant (see Methods). This analysis revealed both well-established immunotherapy response-associated cytokines and previously unrecognized cytokines linked to immunotherapy response.

We identified cytokines that were positively associated with immunotherapy response broadly across multiple cancer types (Figure 5A). These include the IFNγ-chemokine associated gene program [28]. The downstream of IFNγ-chemokine axis is known to create a highly inflammatory and cytotoxic microenvironment [29,30] via (i) gene programs responsible for enhanced chemokine-mediated immune recruitment (CXCL9, CXCL10, CXCL11, CCL4, XCL1, XCL2) [31,32], (ii) increased antigen presentation (B2M) [33], (iii) dendritic cell priming (FLT3LG, CD40LG) [34] which lead to (iv) cytotoxic lymphocyte effector function (GZMB, FASLG) [35] in the TME, consistent with favorable immunotherapy response. We also highlight several potential biomarkers and therapeutic targets revealed by TxCyto such as FLT3LG [36], IL-15 [37], and EBI3 [38], which are associated with favorable response to ICB in multiple cancer types. The complete list of cytokine genes significantly associated with response to ICB therapy across multiple cancer cohorts is provided in Supplementary table 8. These candidate cytokines represent promising biomarkers and potential therapeutic targets that warrant further experimental validation to elucidate their functional roles in regulating anti-tumor immune responses.

**Figure 5.**
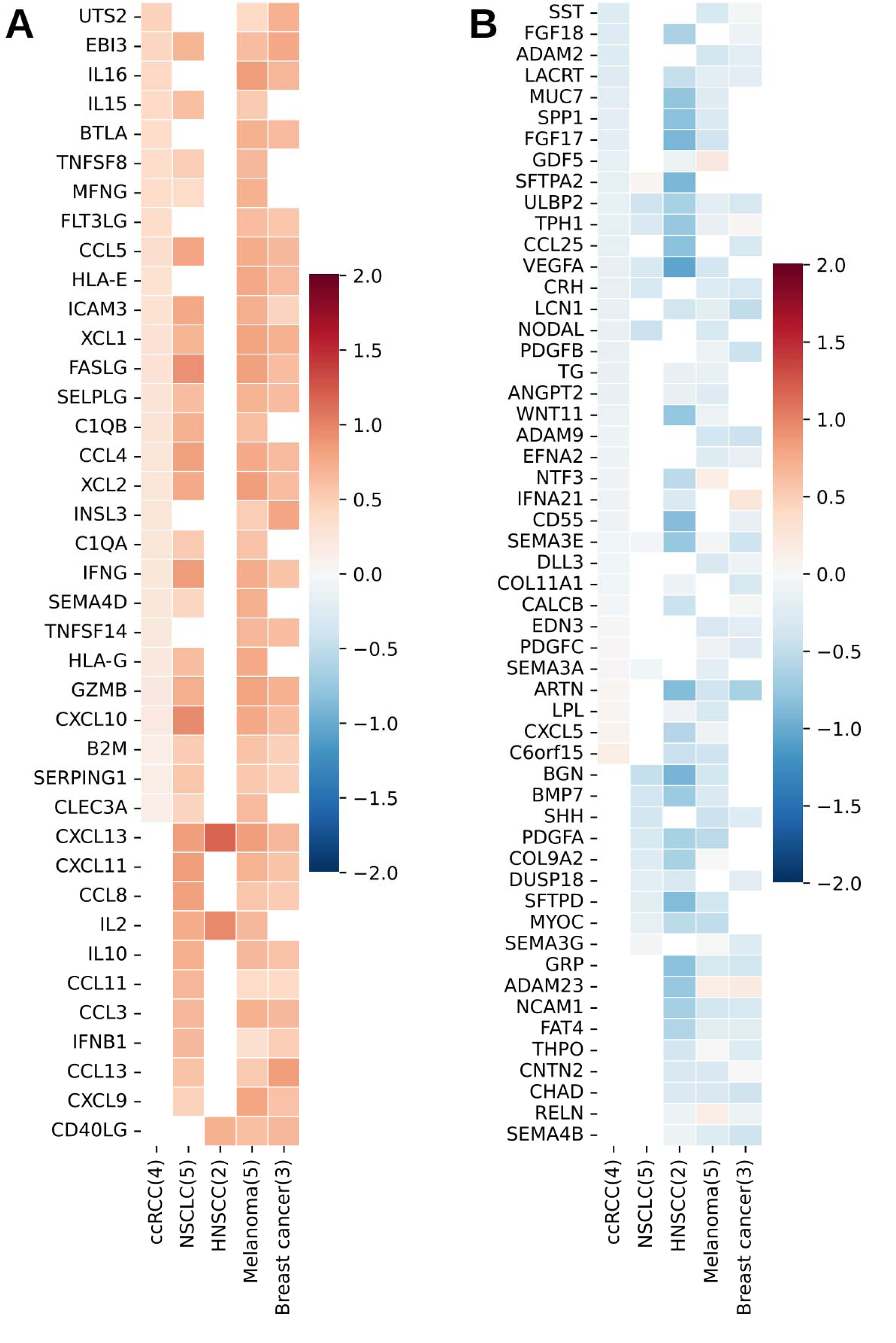
A) The heatmap shows cytokines positively associated with immunotherapy responses across cancer types. X-axis shows tumor types with number of cohorts. Y-axis show cytokines significantly appeared in the majority of cohorts of those cancer types. The color of the heatmap shows the average effect size (treatment-control). B) Similarly, the heatmap shows cytokines negatively associated with immunotherapy responses across cancer types.

In contrast to responder-associated cytokines, which converged on a conserved immune activation program characterized by IFNγ signaling, chemokine-mediated lymphocyte recruitment, and cytotoxic effector function across multiple cancer types, genes associated with non-response displayed substantially greater heterogeneity. These genes largely reflected tumor type-specific and microenvironment-specific programs, including extracellular matrix remodeling, angiogenesis, developmental signaling, and neuroendocrine differentiation. In clear cell renal cell carcinoma (ccRCC) cytokines associated with non-response include those affecting angiogenic and abnormal vasculature (VEGFA, ANGPT2) [39,40] and extracellular matrix remodeling (COL11A1) [41] (Figure 5B), reflecting tumor-extrinsic microenvironmental effects. These findings are consistent with the previously established role of hypoxia-VEGF signaling and vascular remodeling in shaping anti-tumor immune response [39]. In non-small cell lung cancer (NSCLC), the neuroendocrine differentiation (CRH,TPH1) [42,43] and cellular plasticity (SEMA3A, SHH) [44] related cytokines drive the resistance to immunotherapy. Notably, SHH implicates hedgehog signaling, a developmental pathway linked to stemness and therapeutic resistance [45], whereas BGN suggests extracellular matrix remodeling and potential immune exclusion [46]. Again, in head and neck cancer (HNSCC), the cytokines associated with vascular reprogram (VEGFA), cellular plasticity (WNT11, BMP7) [47] are enriched in non-responding samples. In melanoma, cytokines associated with angiogenesis (VEGFA, ANGPT2) [48,49] extracellular matrix remodeling (BGN) [46], activation of developmental pathways (WNT11, SHH, BMP7, NODAL) [47,50], and neural crest-like (NCAM1, NTF3) [51,52] programs were enriched in non-responders (Figure 5B). In breast cancer the identified resistance-associated cytokines are extracellular matrix and stromal remodeling factors COL11A1 [53], neurotrophic signaling such as ARTN [54], which leads to CD8 T-cell exhaustion. Together, these observations suggest that mechanisms of immunotherapy resistance may be highly tissue specific.

Taken together, our results reveal cytokines associated with immunotherapy response across multiple cancer types. It also suggests that, while the cytokine activity associated with immunotherapy response are conserved across cancer types, those associated with resistance may reflect tissue-specific biological mechanisms.

### Spatially resolved tumor-secretome activity identifies tumor niches associated with therapy response

Tumor niche is conventionally defined as a spatial neighborhood in tumor slide with distinct cellular composition. Spatial niches in tumors have been associated with clinical outcome [55]. Here we assessed the extent to which variable niches in spatial transcriptomic data are reflected in spatial cytokine activity profiles and whether the estimated spatially resolved cytokine profiles are predictive of therapy response. We curated two spatially profiled (10X Genomics Visium) cancer cohorts: i) high grade serous ovarian cancer comprising 12 patient samples (6 excellent and 6 poor response to neoadjuvant chemotherapy (NACT) [56] ii) hepatocellular carcinoma (HCC) comprising 7 patient samples (4 responders and 3 non-responders to neoadjuvant cabozantinib and nivolumab therapy) [57] (see Methods). We predicted tumor-secretome activity at each spot using the pre-trained TxCyto model based on the locally smoothed spatial transcriptome in HCC. In the Ovarian cancer cohort, the spatial coordinates were not provided and therefore, we relied on the transcriptome of the individual spots alone without local spatial smoothing to predict the cytokine activity (see Methods).

We used the inferred tumor-secretome activity to classify spots into responders and non-responder to therapy in each cohort. We train a logistic regression classifier, where the secretome activity scores across 645 cytokines at each spot are used as input features. We computed the AUC to assess the accuracy of prediction of spots into responder and non-responders (see Methods). In the HCC cohort, based on leave-one-sample-out strategy (Methods), the spot-level classifier achieved an AUC of 0.75 for distinguishing responder from non-responder regions (Figure 6A). Although the number of donors is very small, we further aggregated spot-level predictions by averaging the responder probabilities across all spots within a slide (model trained on other slides) to derive sample-level predictions; the sample-level analysis yielded an AUC of 0.66. Figure 6C shows the spot-level predicted response probability for the four responder and three non-responder slides. While the majority of spots in responders have higher response probability (first column in Figure 6C), a subset of spots in responder samples (Figure 6C, HCC1R, indicated by black arrow) exhibit higher non-responder probabilities, suggesting the presence of spatially restricted resistant microenvironments corroborated by the enrichment of Exhausted T cell score (Figure 6C, HCC1R, column 5). Similarly in the slide HCC3R, the localized higher non-responding probability is associated with lower infiltration of CD8 T cells (Figure 6C, HCC3R, column 2). The non-responding slides (HCC7NR) have lower enrichment of CD8 T cells and cytotoxic scores (Figure 6C, HCC7NR). Similar to Liver cancer, in the ovarian cancer cohort, the spot-level classifier achieved an AUC of 0.77 for predicting chemotherapy response (Figure 6B), and a sample-level AUC of 0.83. Collectively, these results show that TxCyto-inferred spatial tumor-secretome activity can identify spatial niches in the tumor microenvironment associated with therapeutic response and resistance with good accuracy. Simultaneously, it also provides a framework for characterizing intratumoral heterogeneity based on a comprehensive spatial secretome profile.

**Figure 6.**
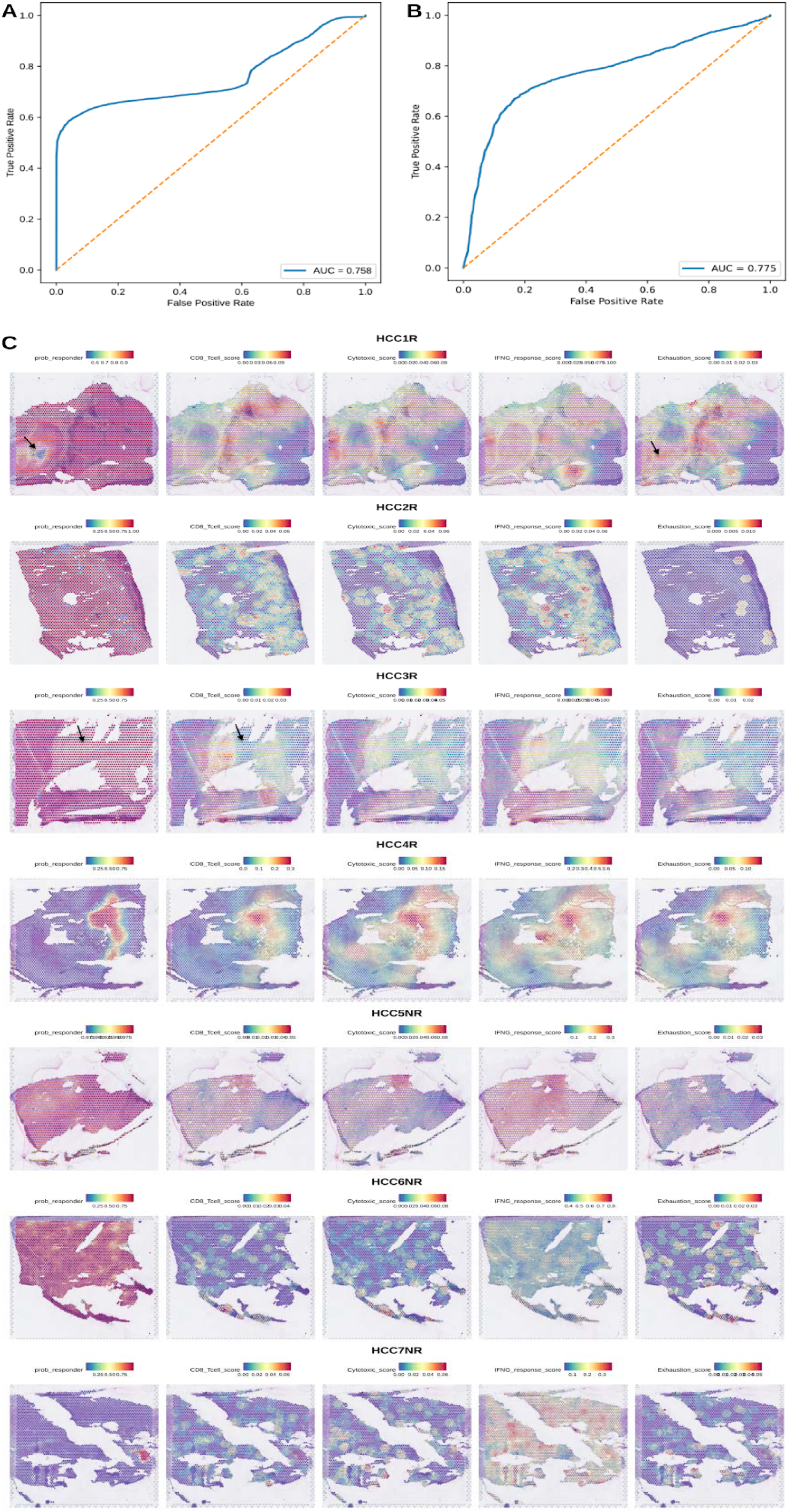
A) The AUROC curve shows the performance of the spot level classifier in distinguishing the response like regions from non-response like regions of novel immunotherapy combination therapies in HCC cohort, based on leave-one-sample-out strategy. B) Similarly in the OV cancer cohort, the AUROC plot shows the performance of the spot-level classifier in distinguishing responder from non-responder regions of neoadjuvant chemotherapy. Both the classifiers use the TxCyto inferred cytokine activity at each spot. C) The plot showing the response probability along with CD8 T-cells, Cytotoxicity, IFN-gamma response and T-cell exhaustion signature score at each spot across all seven HCC spatial transcriptomics slides.

## Discussion

We have presented TxCyto, a deep learning tool to estimate cytokine activity in a transcriptomic sample. TxCyto substantially expands the set of cytokines and other tumor secreted proteins (645 compared to 43 in CytoSig [13], whose activity can be predicted with reasonable accuracy in a bulk transcriptome sample. We have extensively validated TxCyto in multiple independent experimental datasets. TxCyto-predicted cytokine activity tracks the activity of expected downstream processes, demonstrating biological grounding. We have demonstrated the utility of TxCyto in a variety of contexts including identification of cytokines associated with immunotherapy response across multiple cohorts and cancer types, as well prediction of immunotherapy response in ovarian and hepatocellular carcinoma from spatial transcriptomics data, rivaling state of the art. Thus, TxCyto represents a significant advance in translational research.

TxCyto approximates cytokine activity by its global mRNA level. This choice is obviously prompted by data availability. While protein level may be a better approximation of cytokine activity, it is still an approximation, since cytokine activity depends not only on its post-translational modifications, but also on functional interactions [65]. Thus, training a model based on cytokine activity is currently not feasible. We have however shown that the mRNA-based model performs reasonably well in predicting cytokine protein levels in independent cohort, and surprisingly, in some cases, the predicted cytokine level exhibit greater correlation with the protein level of the cytokine than the observed mRNA does, suggesting that the model captures the transcriptional phenocopy of the cytokine activity.

Cytokines are secreted in the TME by multiple cell types and affect the transcriptome of multiple cell types. Thus a faithful measurement of transcriptome of all cells in the tissue should represent overall cytokine level or activity in the tissue. While scRNA-seq data may seem like a better avenue for modeling cytokine activity, both theoretical considerations (disassociation biases, missed rare cells, inaccurate approximation of global cytokine level in the tissue) and practical considerations (primarily relatively limited cohorts) prompted us to train TxCyto in TCGA bulk transcriptome cohort. As we have shown, indeed, bulk-based TxCyto is superior to the scRNA-based model. Furthermore, it is interesting to note that the model based on overall transcriptome (TCGA bulk or pseudo-bulked scRNA-seq) is more accurate than models based on individual cellular compartments (deconvolved cell types in TCGA or cell types in scRNA-seq). This is consistent with the fact that cytokines affect multiple cell types and therefore a collective of all cell types better reflects global cytokine activity in the tissue.

Genes form intricate networks of dependencies and any perturbation in a regulatory molecule such as cytokine, transcription factors, or miRNA, will have a predictable impact, direct and indirect, on the global transcriptome of the tissue [66]. It then stands to reason that shifts in global transcriptome reflect activities of regulatory molecules. We have previously applied this premise to build a machine learning model to predict cellular miRNAs from the transcriptome [67]. Here, we develop a more powerful deep learning tool for global cytokine level prediction. It follows that TxCyto represents a general tool that can be trained to estimate the levels of any regulatory molecule from the transcriptomic data, and the tool and the training code base we provide should serve this purpose for the community.

While TxCyto is trained to predict individual cytokines, given the pleiotropy and redundancy among cytokines, ultimately it is the overall cytokine profile in a tissue that would best characterize its phenotype. This was demonstrated in our application to ovarian cancer spatial transcriptomic data, where a machine learning model based on predicted activities of multiple cytokines better predicts the immunotherapy response, both at tile level and whole slide level. Given its generalizability and ease-of-use, TxCyto represents a powerful tool to investigate the role of cytokines in contexts ranging from aging, autoimmune disease, development, and cancer.

## Methods

### Data collection

We obtained the preprocessed TCGA bulk transcriptomic data from the Broad GDAC Firehose repository https://gdac.broadinstitute.org/ [58]. The final gene expression matrix comprising all 9491 unique patient samples across 33 tumor types is given in the Supplementary tables 10. The corresponding deconvolved gene expression data for these TCGA samples were obtained from the Zenodo repository <u>10.5281/ZENODO.4469784</u> associated with the published study [59]. The single cell RNA-seq data to compare the model was obtained from the published repository <u>10.5281/ZENODO.15554080</u> [22]. This dataset comprises 1062 samples across 6 different cell type populations which includes malignant cells and various immune cell populations. The matched RNA-seq and protein expression profiles for ovarian cancer were obtained from [60]. The comprehensive cytokine perturbation in 24 different cell populations is obtained from [15]. The comprehensive tumor secretome dataset is obtained from the repository https://zenodo.org/records/7074291[19]. The cytokine and its matched receptor is curated from multiple sources and provided in the Supplementary table 5. The meta data and processed gene expression for cytokine induction experiments is obtained from [13]. The GEO accession number for cytokine induction is provided in the Supplementary table 2. The gene expression data from human physiological conditions treated with cytokine inhibitors is obtained from its GEO accession code listed in Supplementary table 3. The analysis for the association of receptor level with cytokine activity is performed on an independent dataset obtained from [61]. The immunotherapy related transcriptomics data is provided with its accession code listed in the Supplementary table 6. The Spatial transcriptomics data (10X Genomics Visium) is obtained from [62,63]. All transcriptomic data were normalized on a per-sample basis, preserving the relative expression levels of genes within each sample before being used as input for the model.

### Model construction and performance evaluation

Figure 1 shows the cytokine prediction model pipeline (TxCyto) based on a fully connected neural network (FCNN). FCNNs are well suited for learning complex relationships among variables [64]. By learning hierarchical representations from whole-transcriptome gene expression profiles, the FCNN captures underlying transcriptional patterns that enable more accurate estimation of cytokine activity than conventional linear machine learning models. It estimates the expression of cytokine (c) from the whole-transcriptome gene expression profile as follows:

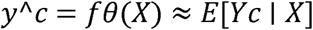

Where,

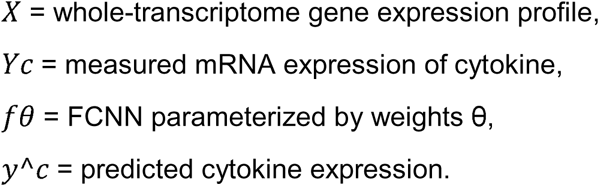

The TCGA samples from 33 distinct tumor types were pooled to construct the training cohort. In total, the dataset comprises 9491 transcriptomics samples with 6075 curated training features. The training features comprise the common genes across the entire training and validation cohorts. We further excluded the 645 tumor secretome from the feature lists, eventually ending up with 6075 genes as the input features to the model. Likewise, cell type-specific transcriptomic profiles from 11 different cell types were integrated to generate the corresponding cell type-specific training matrices. The input features are passed to the model after normalizing it across TCGA samples. The 5-fold cross validation (CV) accuracy is computed using Pearson’s correlation between predicted and actual mRNA expression of the cytokine in the held-out cohort. Next, to evaluate the suitability of the FCNN based model for predicting cytokine activity, we compare its performance with 1) Ridge regression based linear model ii) XGBoost: a decision tree based non-linear model. To be consistent across these three model architectures, we trained and evaluated the models on the same input features and sample matrix. The CV accuracy of cytokine was evaluated using pearson correlation between predicted and observed cytokine level in held out samples. We compared the median CV accuracy of cytokines across these three model architectures. Next, we compared the prediction accuracy of FCNN based models trained on TCGA with those trained on single-cell pseudobulk transcriptomic data. Specifically, CV performance was evaluated for four models: (i) the TCGA bulk transcriptome-based model, (ii) the TCGA cancer cell-specific model, (iii) the single-cell pseudobulk transcriptome-based model, and (iv) the single-cell epithelial pseudobulk transcriptome-based model. Further, we compared the cross-modality predictive performance of the models. The model trained on TCGA bulk transcriptomes was evaluated on single-cell pseudobulk transcriptomic data and vice versa. Both models were independently applied to cytokine-stimulated transcriptomic datasets, and the treatment-induced changes in predicted cytokine activity (delta = treatment − control) were compared.

### Independent validation of the model

We applied TxCyto to transcriptomic datasets generated under human physiological conditions. The pre and post treatment transcriptomics data involving cytokine inhibitors were curated from the GEO accession listed in the Supplementary table 3. For each dataset, we estimated the cytokine activity for each sample at different treatment conditions. Next, we grouped the sample based on either dosage, time point or response to the treatment, depending on the meta data available for that sample cohort. We inferred the change in activity as delta between conditions, calculated as high dosage minus low dosage, later minus earlier time point, or responders minus non-responders, depending on the experimental context. If the delta <0, the model was deemed to be correctly inferring the cytokine activity level. Simultaneously, we compared the estimated cytokine activity score from TxCyto with the score estimated from the CytoSig model for the same group of samples. Next, we applied TxCyto, to single cell RNA-seq data generated from 12 healthy female and male peripheral blood mononuclear cells (PBMCs) donors treated with 90 different cytokines. The cell type-specific pseudobulk transcriptomic profiles were provided for 24 different immune cell populations under both cytokine-treated and control conditions. We removed the immune cell types from the analysis for which all 12 donors’ scRNA seq data is not present. Further, we focused only on those cytokines which have high CV accuracy >0.6. After filtering for the number of donors and cytokines, we ended up with 45 cytokines and 17 immune cell populations from PBMCs. The cytokine activity is estimated pre and post treatment in these 17 different immune cell populations. The cytokine-induced change in cytokine activity (delta = post-treatment − pre-treatment) was estimated for those 45 different cytokines across 17 immune cell populations to assess the model’s predictive accuracy.

Further, we validated the predicted cytokine activity score based on the activity of its canonical downstream pathways. We curated the canonical downstream pathways of 15 important cytokines from the study [25,26]. We computed the pathway score in each sample across different immune cell populations using the ssGSEA pipeline [24]. Pearson’s correlation was computed between the predicted activity of each cytokine and its corresponding downstream signaling pathway across the 17 immune cell populations. The cytokine-downstream pathways pair is provided in the Supplementary Table 4.

Next, we curated 88 unique cytokines and their corresponding receptors from publicly available databases [19], provided in Supplementary Table 5. To evaluate the contribution of receptor expression, we employed the following regression equation to get the β1 coefficient of the receptor is as follows:

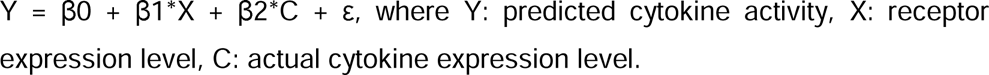

We compared the regression coefficient (β□) of the receptor between the matched cytokine– receptor pairs and randomly paired cytokine–receptor combinations. A one-sided Mann– Whitney U test was performed to determine whether the distribution of the β□coefficients from the actual cytokine–receptor pairs were significantly greater than that of the randomly paired cytokine–receptor.

### Application

Finally, TxCyto was applied to 19 immunotherapy transcriptomic cohorts spanning five different tumor types. Details of the cohorts, including GEO accession numbers, tumor types, and response annotations, are provided in Supplementary table 6 and 7. Predicted cytokine activity was estimated for each sample across all tumor cohorts. The effect size was defined as the difference in cytokine activity between responder and non-responder groups. Statistical significance was assessed using a one-sided Mann–Whitney U test. Two separate tests were performed to identify cytokines enriched in responders (effect size>0) and cytokines enriched in non-responders (effect size<0). P values from both the directions were adjusted for multiple testing using Benjamini-Hochberg procedure. An FDR threshold of <0.25 was used to identify cytokines associated with response or resistance to immunotherapy within each cohort. The consistency of these significant cytokines was then evaluated across cohorts from the same tumor type. Only cytokines showing a consistent direction of association were retained in the final prioritized list. Specifically, consistency was defined as three or more cohorts for clear cell renal cell carcinoma(ccRCC), Non-small cell lung cancer (NSCLC), Melanoma and two or more cohorts for head and neck squamous cell carcinoma (HNSCC) and Breast cancer. Finally, the list of cytokines positively and negatively associated with response and resistance across tumor tissue types are provided in the Supplementary table 8 and shown in Figure 5.

Next, two spatial transcriptomic cohorts (10x Genomics Visium) for ovarian cancer and hepatocellular carcinoma (HCC) respectively were obtained from [56,57]. HCC cohorts comprise seven spatial transcriptomic slides with four responders and three non-responders to novel immunotherapy combination therapies (neoadjuvant cabozantinib and nivolumab). Similarly, Ovarian cancer consists of twelve spatial transcriptomics slides with six responders and six non-responders to neoadjuvant chemotherapy (NACT). First, a locally smoothed gene expression profile was computed for each spot in the HCC spatial transcriptomics cohort. The HCC data was generated using standard 10X genomics Visium platform, in which the spots are arranged on a hexagonal grid. Gene expression was smoothed by averaging the expression profiles of neighboring spots within a radius of 120 pixels, corresponding approximately two concentric rings around the central spot. Depending on the spot location within the tissue, this neighborhood comprised an average of 16 spots. In contrast, the ovarian cancer spatial transcriptomics dataset did not include spot-level spatial coordinates. Therefore, cytokine activity was predicted directly from the preprocessed gene expression profile of each individual spot without additional spatial smoothing. Then, all the 645 predicted cytokine activity scores were used to classify the spots into responders and non-responders using logistic regression in a leave-one-slide-out cross validation strategy. A similar approach is applied to classify the spots into responder and non-responder in ovarian cancer cohorts. The AUROC for both the data cohorts is provided in Figure 6. Next, we calculated CD8 T-cells, Cytotoxicity, IFN-gamma response and T-cell exhaustion signature score at each spatial spot across all seven HCC tissue slides. The signature genes for each of these four markers are provided in the Supplementary table 9. Further, in both cohorts, a slide-level AUROC is computed by averaging the predicted probability across all the spots within a slide to obtain a single slide level prediction. The spot level and sample level prediction probability of responder and non-responders for ovarian cancer data cohort is provided in the Supplementary table 11. Similarly, the spot level and sample level prediction probability in HCC tumor cohort is provided in Supplementary table 12.

## Data and code availability

All the codes used for the analysis of the manuscript, intermediate files, notebooks, supplementary tables along with instructions to reproduce the results are publicly available at https://github.com/Rahulncbs/TxCyto.

## Acknowledgements

This work used the computational resources of the NIH HPC Biowulf cluster. We thank Arashdeep Singh, Vishaka Gopalan and Gulden Olgun for useful discussion and valuable feedback. We also thank Sumit Mukherjee and Lipika Ray for providing the immunotherapy dataset used in this study. We are grateful to Beibei Ru for directing us to single cell transcriptomics data used in the analysis. We used OpenAI’s ChatGpt (GPT-5.6 Sol) to assist with editing text in the methods sections of the manuscript.

## Contributions

R.K.: Study design, data curation, software, formal analysis, investigation, visualization, methodology, writing–original draft, writing–review and editing. S.L.: formal analysis. P.J.: supervision, writing–review. S.H.: Supervision, conceptualization, study design, funding acquisition, investigation, project administration, writing–original draft, writing–review and editing.

## Supplementary Figures

**Supplementary Figure 1.**
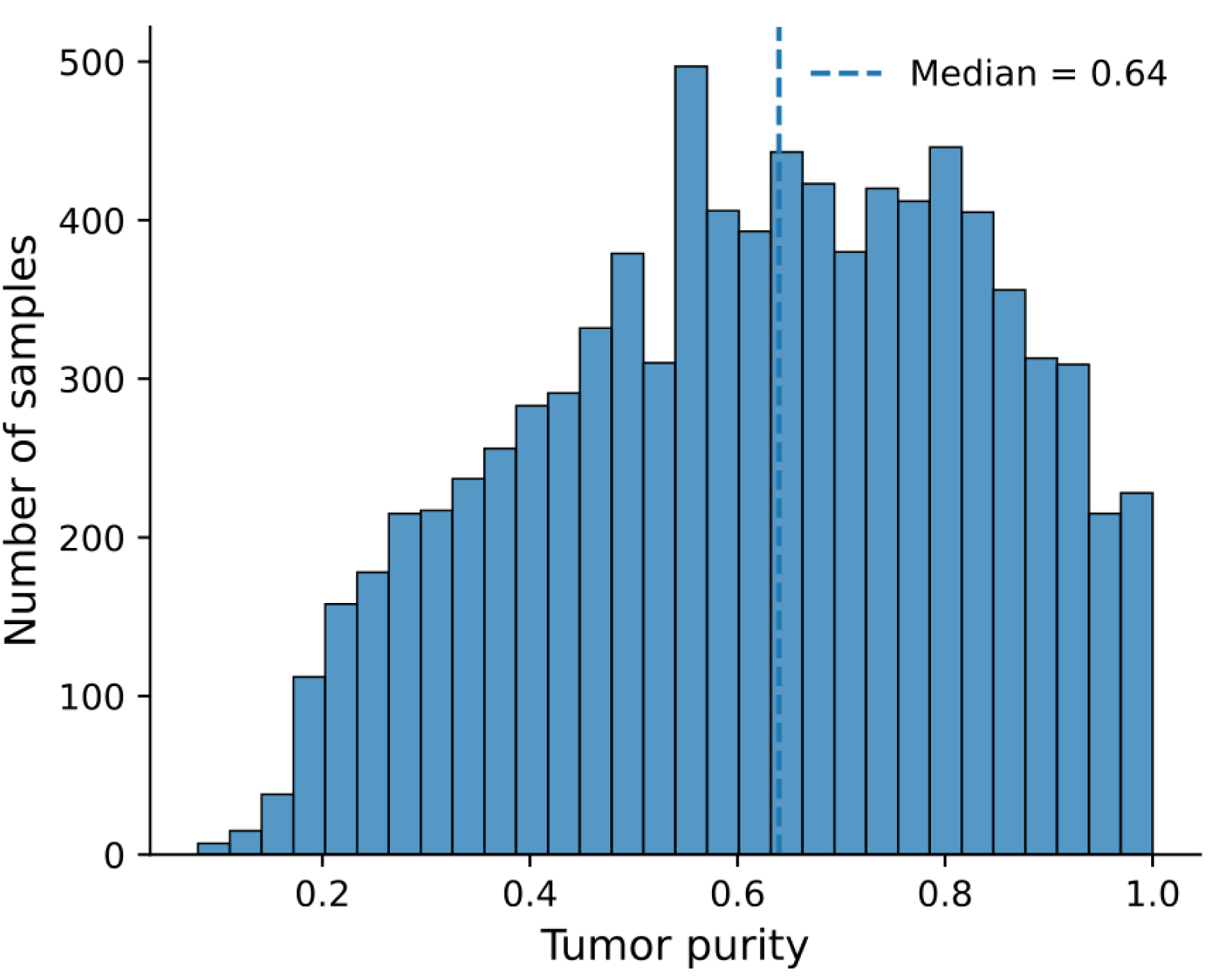
The distribution of the purity score of TCGA tumor samples having median purity score∼0.65.

**Supplementary Figure 2.**
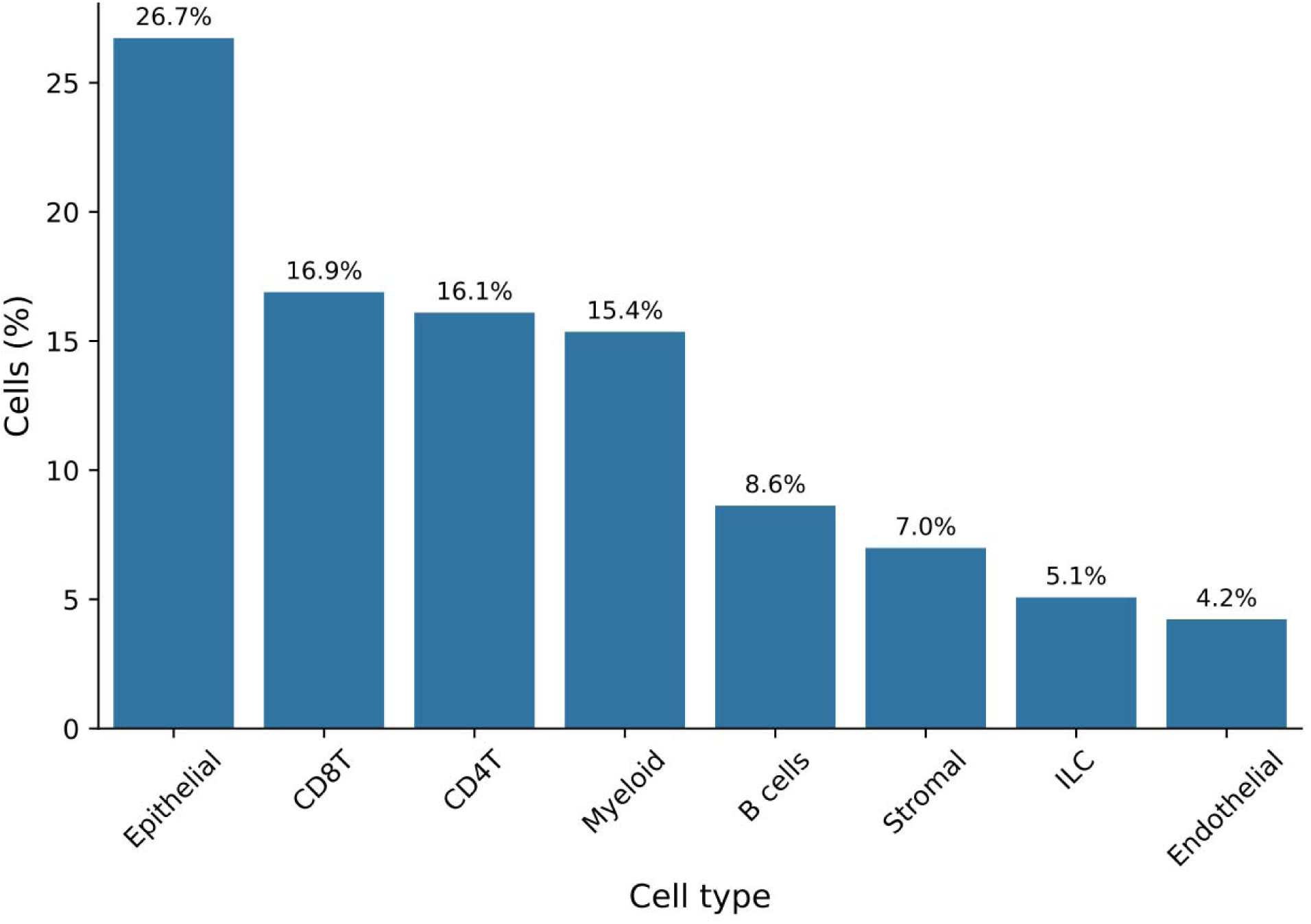
The plot shows the proportion of different cell types across 1062 single-cell RNA-seq samples.

**Supplementary Figure 3.**
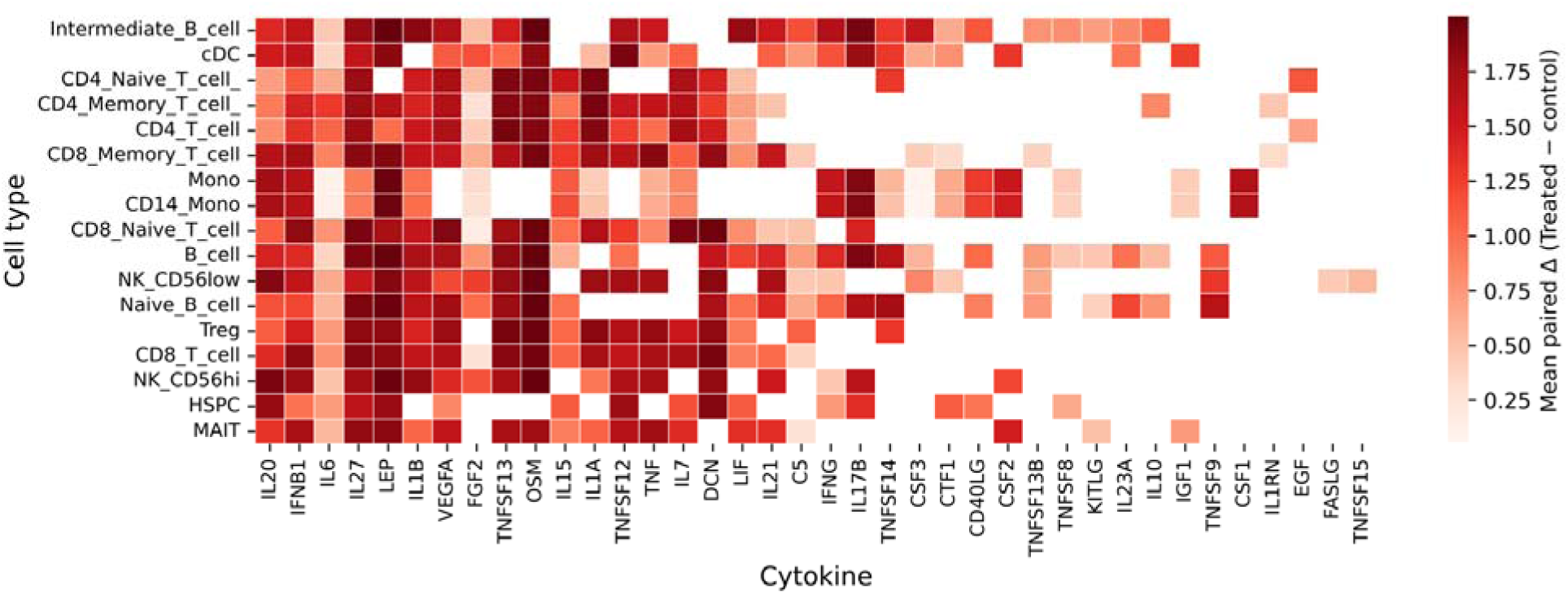
The heatmap shows the significantly elevated cytokines (p-value <0.05) across different immune cell populations of treated PBMC samples.

